# A PCR-RFLP method for the detection of CRISPR-induced indels

**DOI:** 10.1101/2023.04.09.535589

**Authors:** Lydia Angelopoulou, Electra Stylianopoulou, Konstantinos Tegopoulos, Ioanna Farmakioti, Maria E. Grigoriou, George Skavdis

## Abstract

CRISPR-based technologies have revolutionised genome editing and are widely used for knocking out genes in cell lines and organisms. From a practical perspective, a critical factor that largely influences the successful outcome of CRISPR gene knockout experiments is the reliable and fast identification of fully mutated cells carrying exclusively null alleles of the target gene. Here we describe a novel strategy based on the well-documented reliability and simplicity of the classical PCR-Restriction Fragment Length Polymorphism (RFLP), which allows the assessment of the editing efficiency in pools of edited cells and the effective identification of cell clones that carry exclusively mutated alleles. This fast and cost-effective method, named PIM-RFLP (PCR Induced Mutagenesis-RFLP), is executed in two steps. In the first step, the editing target is amplified by a set of mutagenic primers that create a restriction enzyme degenerate cleavage site in the amplification product of the wild type allele. As a proof of principle, we chose the XcmI restriction site because it is especially suitable since it has the particularity of containing nine centrally placed non-specific nucleotides. This gives great flexibility in the mutagenic primers design and allows for efficient execution of the mutagenic PCR. In the second step, the evaluation of the editing efficiency in pools of edited cells or the identification of fully mutated single-cell derived clones is achieved following the standard procedure for any PCR-RFLP assay: digestion of the PCR products and analysis of the restriction fragments in an agarose gel.

**GRAPHICAL ABSTRACT:** 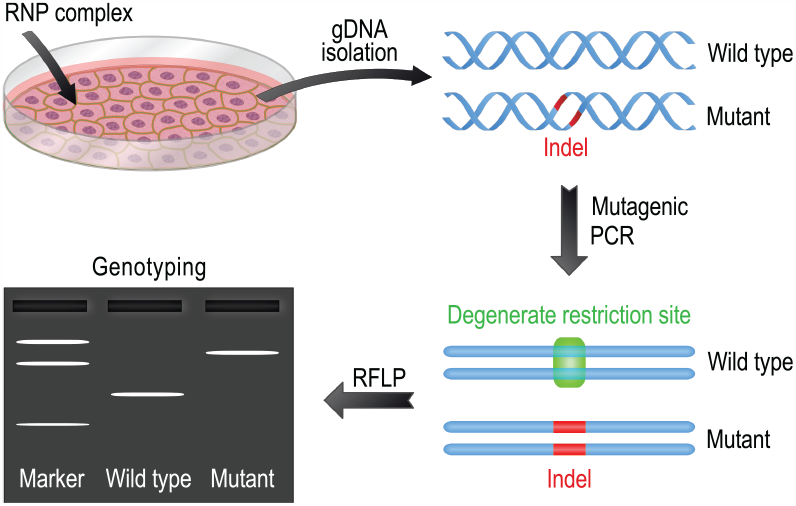

## INTRODUCTION

CRISPR genome editing technology is currently the most widely used approach for knocking out genes. In a nutshell, this strategy is based on using a predesigned guide RNA (gRNA) to recruit an appropriate nuclease (usually SpCas9) to a specific gene sequence. The nuclease introduces a DNA double-strand break (DSB), which is often repaired by an error-prone mechanism that generates mutated alleles of the target gene (Knott & Doudna, 2018; Komor *et al*, 2017). The majority of these alleles are characterised by the presence of relatively small indels around the repaired DSB and are functionally null (Chakrabarti *et al*, 2019; Chen *et al*, 2019; Allen *et al*, 2019).

Next-generation sequencing (NGS) of PCR products encompassing the DSB is considered the gold standard allele profiling method for analysing CRISPR-mediated knockout experiments, because it allows the identification and quantitation of practically every allele present in a pool or clone of edited cells. However, the cost of this approach is often prohibitive, and since NGS is not an in-house technique for most laboratories, the turnaround time from samples to results can be considerably long. These practical limitations have fuelled the development of a plethora of alternative methods aiming to analyse CRISPR editing results (Bennett *et al*, 2020).

One of the first methods that was widely used, and is still a popular choice, is the T7 endonuclease 1 (T7E1) mismatch cleavage assay, which is based on the ability of T7 endonuclease 1 to digest heteroduplex DNA molecules produced from the denaturation and renaturation of PCR amplicons encompassing the DSB. These heteroduplexes are formed when partially complementary DNA strands from different alleles anneal to each other during the renaturation step. An inherent limitation of T7E1 assay is caused by the fact that the amount of heteroduplex DNA formed does not always reflect the amount of editing. For example, in highly edited samples with a single prevailing mutant allele, mostly homoduplexes are formed. Hence, in pools of edited cells, T7E1 assay greatly underestimates the editing efficiency when the commonest indel frequency is high (Sentmanat *et al*, 2018). Furthermore, singlebase loops can be resistant to digestion by T7E1 (Yang *et al*, 2015) and thus, 1 bp indels, which are the most common frameshift mutations obtained from CRISPR knockout experiments (Chakrabarti *et al*, 2019; Allen *et al*, 2019; Chen *et al*, 2019), can remain undetected. Therefore, T7E1 assay can fail to correctly genotype single-cell derived clones containing mutated alleles with 1 bp indels (Yang *et al*, 2015).

A new generation of allele profiling software-based methods have been gaining popularity during the last few years. They all use Sanger sequencing deconvolution algorithms, which are capable of deciphering complex Sanger traces obtained from the sequencing of a heterogeneous amplicon population produced by PCR amplification of the region surrounding the DSB (Brinkman *et al*, 2014; Bloh *et al*, 2021; Conant *et al*, 2022). These methods are cost-effective and more reliable than the T7E1 assay (Sentmanat *et al*, 2018). They also have the advantage over the T7E1 assay of providing sequencing information for the detected alleles. However, they can also have a relatively long turnaround time, especially in the case of laboratories that do not have in-house Sanger sequencing services.

Here we describe a new PCR-RFLP approach for the detection of CRISPR-induced indels. We named this method PCR Induced Mutagenesis Restriction Fragment Length Polymorphism (abbreviated as PIM-RFLP) because it is based on the creation of a degenerate restriction site on the PCR product of the wild type (unedited) allele. The XcmI restriction enzyme is especially suitable for PIM-RFLP for several reasons. Firstly, and more importantly, its recognition site (5′-CCANNNNN/NNNNTGG-3′, abbreviated thereafter as 5′-CCAN_9_TGG-3′) contains nine centrally placed random nucleotides. This allows the design of mutagenic primers with only few and distantly placed from the 3′ end mismatches, since the position of the mismatches is not fixed. Secondly, the XcmI cleavage site contains the dinucleotide GG. This simplifies the design of the mutagenic PCR, as SpCas9 (the most commonly used programmable endonuclease) recognises the PAM sequence 5′-NGG-3′. Thirdly, since XcmI is a 6-cutter restriction enzyme, its presence is relatively rare. Based on these properties of XcmI, we chose this enzyme to provide a proof of principle for PIM-RFLP. PIM-RFLP starts with the isolation of genomic DNA (gDNA) from the samples of interest, which then serves as PCR template using a pair of mutagenic primers flanking the DSB. As it can be seen in Figure 1, through this procedure the wild type allele is amplified to produce an amplicon that carries an XcmI recognition site. In contrast, alleles containing deletions extending up to 1 bp (on either or both sides of the DSB), as well as insertions, produce amplicons that lack the XcmI site, while alleles containing larger deletions are generally not amplified. The amplified edited alleles can be distinguished from the wild type by XcmI digest followed by agarose gel electrophoresis. We have extensively compared PIM-RFLP with NGS and ICE, a popular Sanger sequencing deconvolution method developed by Synthego Corporation (Conant *et al*, 2022). Here we present data indicating that PIM-RFLP provides highly dependable results and can outperform ICE. Since PIM-RFLP is a reliable, low-cost method with minimal instrumentation requirements and can provide same-day results, it makes a very attractive choice for the analysis of CRISPR knockout experiments.

**Figure 1.**
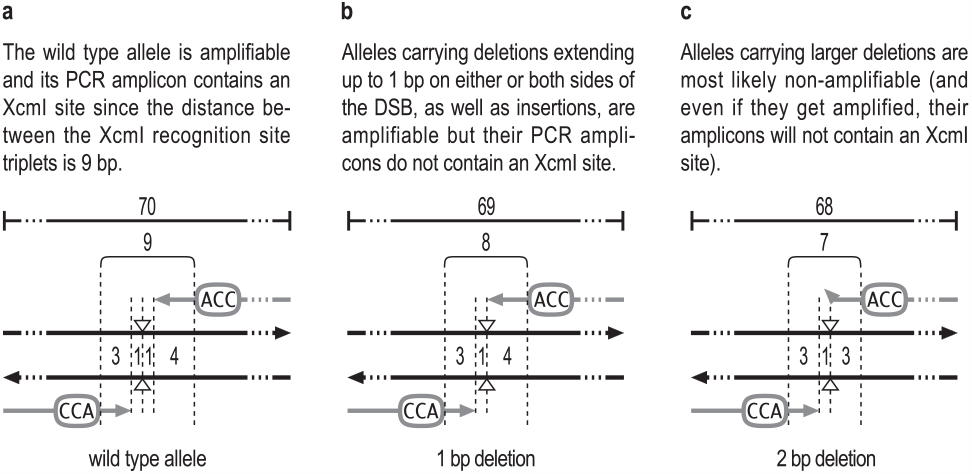
Schematic representation of PIM-RFLP. The template (black arrows) is PCR amplified using a pair of mutagenic primers (gray arrows, on which the specific nucleotide triplets of the XcmI recognition site are depicted) to produce amplicons encompassing the DSB position, which is indicated by a pair of open arrowheads facing each other. All numbers denote base pairs. Brackets indicate the distance between the XcmI recognition site triplets. **(a)** The primers are designed to amplify the wild type allele, producing a 70 bp amplicon that contains an XcmI recognition site (5′-CCAN_9_TGG-3′). Thus, in this example, the positions −4 to −6 (relatively to the 3′-end) of the forward primer and the positions −5 to −7 of the reverse primer are occupied by the triplet 5′-CCA-3′. However, in general, since the positions of the mismatches are not fixed, any design that places these two triplets 9 nucleotides apart can allow efficient amplification as long as the number of primer-template mismatches (especially those that are closer to the 3′ end of the primers) is kept to a minimum. The 3′ end of both primers is located 1 nucleotide before the DSB. **(b)** Mutated alleles with deletions extending up to 1 bp on either or both sides of the DSB, as well as insertions of any size, are also amplifiable but carry the sequence 5′-CCAN_#9_TGG-3′, which is not recognised by XcmI. **(c)** Mutated alleles carrying larger deletions (like the 2 bp deletion depicted here) are most likely non-amplifiable because of mismatches between their sequence and the 3′ end of one or both PCR primers (the bended arrow in the reverse primer indicates the presence of a 3′mismatch). Depending on the target locus sequence, a minority of alleles of this category can be amplifiable, however their amplicons will carry the sequence 5′-CCAN_<9_TGG-3′, which is not digested by XcmI.

## MATERIALS AND METHODS

### Cell culture and transfection

The Neuro-2a cell line (ATCC CCL131) was cultured at 37 °C and 5% CO_2_ in DMEM high glucose (Pan Biotech), supplemented with 10% v/v FBS (Gibco), 1 mM Penicillin-Streptomycin (Pan Biotech) and 2 mM L-Glutamine (Pan Biotech). For the CRISPR editing experiments, 20,000 cells were seeded in a 24-well plate and after 48 hours they were transfected with a ribonucleoprotein complex containing Alt-R S.p. HiFi Cas9 Nuclease V3 (Integrated DNA Technologies IDT) and either a sgRNA that targets the *Abracl* gene (IDT) or a crRNA:tracrRNA that targets the *Hprt* gene (mouse Alt-R CRISPR-Cas9 Control Kit, IDT). Transfection was performed using Lipofectamine CRISPRMAX Cas9 Transfection Reagent (Invitrogen) according to the manufacturer’s instructions. The sequences targeted by the gRNAs are presented in Supplementary Table S1. Two days after transfection, cells were trypsinized and harvested. The collected pools of cells were used for the isolation of single-cell clones. For clonal selection, cells were plated into a 10 cm culture dish at a density of 500 cells per plate and when individual colonies were formed they were picked and expanded. Alternatively, the cells were seeded into 96-well plates at a density of one cell per well.

### Genomic DNA isolation

Liver gDNA was extracted using the NucleoSpin Tissue kit (Macherey-Nagel) according to the manufacturer’s instructions. For Neuro-2A gDNA extraction, briefly, the cells were lysed in a buffer containing 10 mM Tris HCl (pH 8.0), 10 mM EDTA (pH 8.0), 0.08 % SDS, 10 mM NaCl, 400 μg/ml Proteinase K and 100 μg/ml RNase A. Lysis was achieved by 1 h incubation at 55 °C under shaking. After lysis the gDNA was isopropanol precipitated, the pellet was washed twice with 70% EtOH, air-dried and dissolved in water. The gDNA concentration was determined using the NanoDrop 2000c spectrophotometer (Thermo Fisher Scientific).

### PIM-RFLP

Mutagenic PCR was carried out in 20 μl reactions, using 10 μl of KAPA SYBR FAST qPCR Master Mix (2X) ABI Prism (Sigma-Aldrich), which contains a Taq DNA polymerase that lacks proofreading activity, 0.1 μM of each primer and 50 ng of gDNA. All mutagenic primers were polyacrylamide gel (PAGE) purified and their sequence is presented in Supplementary Table S2. Amplification was performed in a StepOne Real-Time PCR System (Applied Biosystems) under the following conditions: one step of 95 °C for 2 min was followed by 32-34 cycles of 95 °C for 10 sec and 60 °C for 10 sec. The concentration of the PCR products was normalised according to their final ΔRn. Subsequently, 10-15 ng of each PCR product were digested at 37 °C for 60 min with 2.5 units of XcmI (New England Biolabs) in 15 μl reactions, containing 1.5 μl of NEBuffer r2.1 (New England Biolabs). Finally, electrophoresis was performed in 4% (w/v) agarose gels prepared with UltraPure Agarose (Invitrogen) and prestained with GelRed Nucleic Acid Gel Stain (Biotium). To estimate the efficiency of editing in pools of edited cells, the PCR reactions were performed in triplicate and average ΔRn values were obtained from the last PCR cycle in which both the edited and the unedited samples were still in the exponential phase.

### Next-generation sequencing

For the *Abracl* locus a target region of 198 bp was amplified using the set of primers FA/RA, while for the *Hprt* locus a target region of 200 bp was amplified using the set of primers FH/RH (Supplementary Table S2, see also Supplementary Figure S1). PCR products were purified using magnetic beads (NucleoMag NGS Clean-up and Size Select, Macherey-Nagel) according to the manufacturer’s instructions in a DNA to beads volume ratio of 1:1.8. The concentration of the purified PCR products was determined in a Qubit 4 Fluorometer (Thermo Fischer Scientific) using Qubit dsDNA HS Assay Kit (Invitrogen) following the manufacturer’s guidelines. The libraries were prepared from 100 ng of purified PCR products using the Ion Plus Fragment Library Kit (Thermo Fischer Scientific) as recommended by the manufacturer and they were purified using the NucleoMag magnetic beads in a DNA to beads volume ratio of 1:1.4. The libraries were then quantified by real-time PCR using the Ion Universal Library Quantitation kit (Thermo Fischer Scientific) according to the manufacturer’s instructions and template preparation was carried out on an Ion Chef System (Thermo Fischer Scientific) following the manufacturer’s guidelines. Finally, sequencing was performed by Ion Torrent GeneStudio S5 (Thermo Fischer Scientific). The obtained data were analysed using CRISPResso2 (Clement *et al*, 2019) under default parameters for the *Hprt*, while for the *Abracl* the minimum single bp quality (phred33 scale) was set higher (>10). For clone genotyping, alleles appearing at <2% frequency were omitted from the analysis and the allele frequencies were recalculated based on the total number of the remaining reads.

### Sanger Sequencing and ICE

For the *Abracl* locus a target region of 688 bp was amplified using the F1A/R1A pair of primers, while for the *Hprt* locus a target region of 647 bp was amplified using the F1H/R1H pair of primers (Supplementary Table S2, see also Supplementary Figure S1). PCR products were purified using the NucleoMag magnetic beads (in a DNA to beads volume ratio of 1:1.8). About 175 ng of purified PCR fragments were Sanger sequenced by StarSEQ GmbH (Mainz, Germany). The sequencing traces were analysed using the online web tool ICE (https://ice.synthego.com/).

## RESULTS

### The mutagenic PCR for creating the XcmI site is very efficient

The special feature of the XcmI recognition site to contain nine random nucleotides between two specific nucleotide triplets, gives great flexibility in designing the mutagenic primers, since the positions of the mismatches between the primers and the template are not fixed. This allows the selection of primers with a reduced number of mismatches and without any mismatch located very close to the 3′ end. In order to test if these XcmI properties allow efficient amplification in the mutagenic PCR step, we randomly chose 10 mouse housekeeping genes plus the *Hprt* gene, because it is commonly used as a positive control in CRISPR editing experiments (an appropriate kit, the Alt-R CRISPR-Cas9 Control, is provided by IDT). For this set of genes, we tested if we could generate by mutagenic PCR a diagnostic XcmI recognition site spanning the DSB, which could be created by Cas9 using for the 10 randomly chosen genes the first ranked gRNA among those suggested by the CHOPCHOP web tool (Labun *et al*, 2019) (Supplementary Table S3), and for the *Hprt* gene the gRNA provided by the IDT kit (Supplementary Table S1). All 11 pairs of mutagenic primers were designed to generate a 70 bp amplicon (Figure 2a). The average Tm of the 22 primers was 69 °C while the highest was 79 °C and the lowest was 61 °C (Supplementary Table S2). After a minimal optimisation procedure, mainly consisting of trying two alternative annealing temperatures (60 °C and 65 °C), we found that most primer pairs worked very well in both temperatures and we could achieve very satisfactory amplification results for all 11 PCR reactions after 32 cycles of amplification at 60 °C (Figure 2b). As it is shown in Figure 2b, in all cases, we were able to produce adequate quantities of amplicons that were digestible by the XcmI. In some digests, a faint undigestible band appeared, which we anticipated that it will not interfere with the analysis of pools of CRISPR edited cells and with the identification of fully mutated single-cell clones (see Results and Discussion).

**Figure 2.**
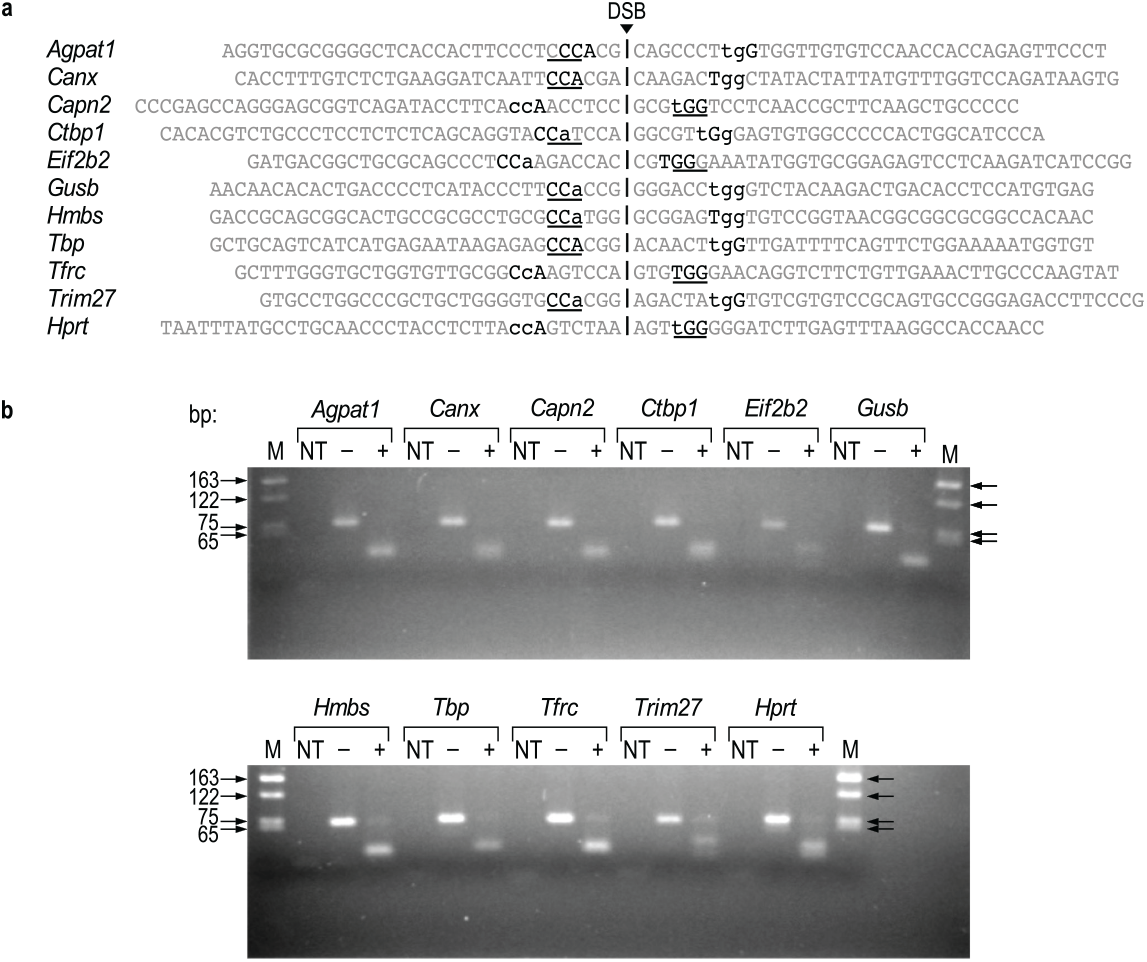
Assessment of the efficiency of the mutagenic PCR used to create an XcmI site **(a)** The 70 bp amplicons (presented in the 5′ to 3′ direction) of the 11 genes tested. Vertical lines indicate the DSB positions. The PAM sequences are underlined. Black letters indicate the specific triplets of the XcmI site. The mutations introduced by the primers are shown in lowercase. The primers extend up to one nucleotide before the DSB. **(b)** Electrophoresis on a 4% (w/v) agarose gel of untreated (–) and treated (+) with XcmI PCR products, obtained using as template mouse liver gDNA. All primer pairs achieved very satisfactory amplification results, and all amplicons were digestible by XcmI. M: DNA marker, NT: Non template control.

### Using PIM-RFLP to evaluate the editing efficiency in pools of edited cells

In PIM-RFLP the wild type alleles are PCR amplified and their amplicons contain an XcmI site. The amplification of mutated alleles is as efficient as that of the wild type (see Figure 1b), or it is hampered (most often is completely abolished) by mismatches between their sequence and the 3′ end of one or both PCR primers (see Figure 1c). Furthermore, the amplicons produced by mutated alleles can be discriminated from their wild type counterparts because they are missing the XcmI site. Thus, when PIM-RFLP is performed in a sample of pooled edited cells using unedited cells as control, the achieved editing is manifested in two ways: (a) the amount of the PCR amplified molecules in the sample of the edited cells is reduced, and/or (b) a fraction of the PCR amplified molecules in the sample of the edited cells is indigestible by XcmI. Thus, the efficiency of the editing can be estimated by quantifying those two effects using the following equation:

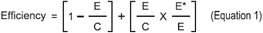

where E and C are the numbers of the amplified DNA molecules in the PCR of the edited and the control (unedited) samples respectively, while E^*^ is the number of the amplified, indel containing (and consequently indigestible by XcmI) molecules in the PCR of the edited sample. The first part of the sum quantifies the non-amplifiable edited alleles, while the second part quantifies the amplifiable edited alleles. The E/C quotient can be calculated through quantitative real-time PCR using the ΔRn values of the edited (E_ΔRn_) and unedited (C_ΔRn_) sample in the exponential phase. The E^*^/E quotient can be calculated by using agarose gel densitometry to measure the intensity of the 70 bp PCR product of the edited sample after (E^*^ = E_Intensity of indigestible_) and before (E = E_Intensity of predigested_) treating the PCR end product with XcmI. Hence, the Equation 1 can be transformed as follows:

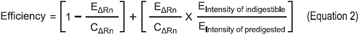

As a proof of principle, using Equation 2, we assessed the editing efficiency in pools of Neuro-2a cells edited at two different loci: the *Abracl* and the *Hprt*. Equal amounts of gDNA from pools of edited and unedited cells were amplified for 34 cycles by real-time PCR using appropriately designed mutagenic primers. The ΔRn values were obtained from cycle 31 for the *Abracl* and from cycle 30 for the *Hprt*, when both the edited and the unedited (control) reactions were still in the exponential phase. To calculate the fraction of the indel containing molecules produced in the edited sample reactions, the corresponding PCR end products were treated with XcmI and electrophoresed in a 4% (w/v) agarose gel (Figure 3). An image of the gel was then analysed with ImageJ software to gather the necessary intensity values. Finally, we assessed the editing efficiency for the *Abracl* and the *Hprt* loci, and we compared the results of PIM-RFLP with those obtained by NGS and by ICE. The results are presented in Table 1. As it can be seen, the editing efficiency values obtained from NGS are closer to those obtained by PIM-RFLP than to those obtained by ICE.

**Table 1:** The CRISPR-editing efficiency in pools of Neuro-2a cells targeted at the *Abracl* and *Hprt* loci evaluated by PIM-RFLP (using the Equation 2), NGS and ICE.

| Gene | $E_{\Delta R_n}$ | $C_{\Delta R_n}$ | $E_{\text{Intensity of indigestible}}$ | $E_{\text{Intensity of predigested}}$ | % Efficiency | | |
| --- | --- | --- | --- | --- | --- | --- | --- |
|  |  |  |  |  | PIM-RFLP | NGS | ICE |
| *Abrac1* | 0.29 | 0.71 | 3,173.01 | 9,568.21 | 72.70 | 68.36 | 49 ($R^2$ 0.93) |
| *Hprt* | 0.41 | 0.61 | 2,179.53 | 8,265.89 | 50.51 | 56.64 | 31 ($R^2$ 0.97) |

**Figure 3.**
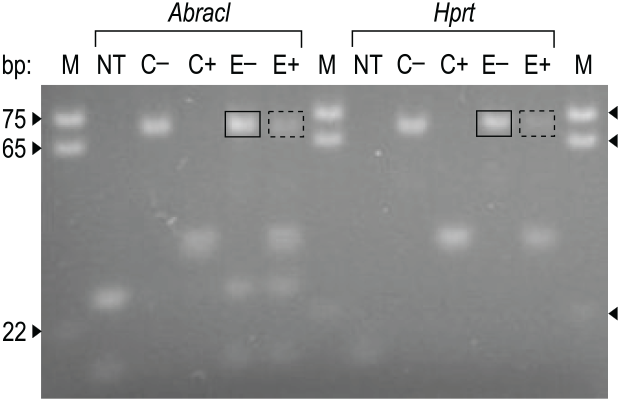
Electrophoresis in a 4% (w/v) agarose gel of untreated (–) and treated (+) with XcmI PCR products obtained using as template gDNA from unedited control cells (C) and cells CRISPR-edited (E) at the *Abracl* and *Hprt* loci. The amplification was performed using mutagenic primers, which generate an XcmI recognition site at the wild type allele amplicon. For each locus the intensity of the 70 bp PCR amplicon produced from the edited sample before (solid line rectangle) and after (dashed line rectangle) the XcmI treatment was measured with the ImageJ software, and the collected values were used for the calculation of the editing efficiency using Equation 2. Although the control reactions produced significantly more product (see ΔRn values in Table 1), here in all lanes approximately the same quantity of PCR products was loaded. M: DNA marker, NT: Non template control.

### Using PIM-RFLP to identify fully mutated single-cell derived clones

We then proceeded picking 23 single-cell derived clones for each of the two edited loci. All 46 individual clones were analysed by PIM-RFLP, NGS and ICE. For PIM-RFLP analysis, 50 ng of gDNA from each clone were amplified for 34 cycles by real-time PCR using the mutagenic primers. Subsequently, all the PCR end products with ΔRn>1 were brought to the same concentration based on their ΔRn values and about 10-15 ng were digested with XcmI and electrophoresed in a 4% (w/v) agarose gel. Potentially negative reactions with ΔRn<1 were treated as having ΔRn = 1. Finally, the clones were genotyped by visual inspection of a gel image (Figure 4) and classified in three categories. When no PCR product was detected (as it happened in all the reactions with ΔRn<1) or the PCR product was completely indigestible by XcmI, the clone was identified as fully mutated and assigned as –. When the PCR product was partially digested by XcmI, the clone was assigned as +/– to indicate the presence of both the wild type allele and at least one mutated amplifiable allele. When the PCR product was fully digested by XcmI, the clone was assigned as +/? because we couldn’t exclude the presence of non-amplifiable edited alleles. It is important to notice that the assigned genotypes denote allelic composition (the presence of wild type and/or mutated alleles) without implying ploidy status (the Neuro-2A cells we used are polyploid having a modal number of 95).

**Figure 4.**
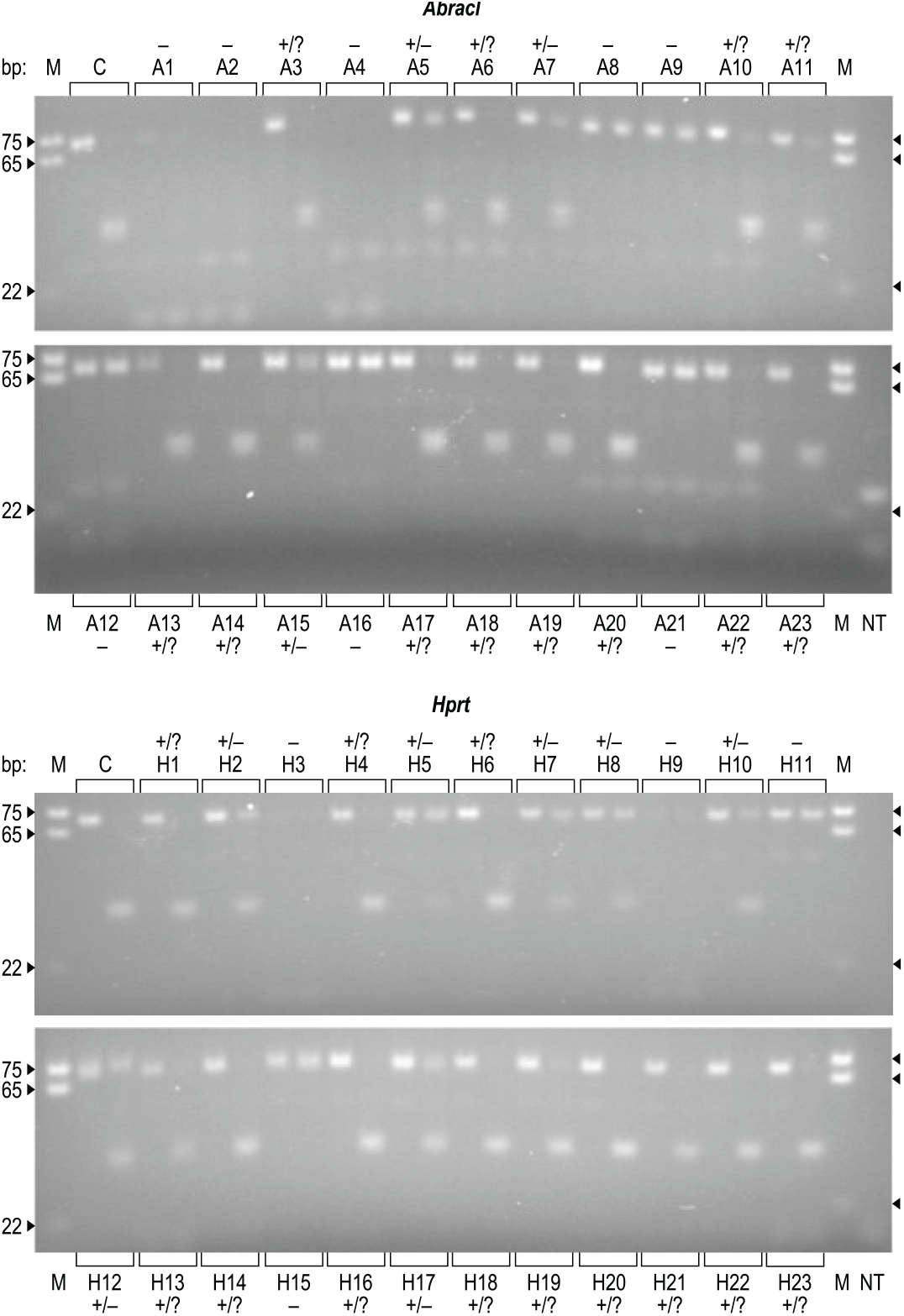
Electrophoresis in a 4% (w/v) agarose gel of PCR products obtained using as template gDNA from single-cell clones derived from pools of Neuro-2a cells CRISPR-edited at the *Abracl* and *Hprt* loci. The amplification was performed using mutagenic primers which generate an XcmI recognition site at the amplicon of the wild type allele. gDNA from unedited Neuro-2a cells was used as control (C). The *Abracl* targeted clones are indicated with an A, while the *Hprt* targeted clones are indicated with an H. The PCR product of each clone was loaded to the gel in a pair of lanes: the left lane contains the amplicon before the XcmI treatment, while in the right lane the amplicon has been treated with XcmI. For each clone the PIM-RFLP assigned allelic composition is shown. Fully mutated clones are indicated by –. Clones containing the wild type and at least one amplifiable mutant allele (revealed by the presence of indigestible PCR product) are assigned as +/–. The potential presence of mutated non-amplifiable alleles is indicated by a question mark and thus, clones that are fully digestible are assigned as +/?. Since a very faint undigested band cannot with certainty be attributed to the presence of indigestible PCR product instead of a partial digest, the clones A10, A11, H13 and H19 are classified as +/? instead of +/–. All the fully mutated clones (–) can be identified unequivocally. M: DNA marker, NT: Non template control.

The results collected by all three methods are presented in Table 2. In most cases, there was a concordance between all methods, however, there were few exceptions. Most importantly, ICE failed to genotype the fully mutated clones A2, A4, A12 and H3, although we used two alternative sequencing primers (Supplementary Figure S1) in an effort to obtain R^2^ values >0.8 which indicate a reliable analysis (Synthego, 2019). This is not surprising since all these clones contain large deletions clearly detected by agarose gel electrophoresis (Figure 5a) and it is known that deletions >40 bases can impact the R^2^ values (Synthego, 2019). The NGS analysis also revealed alleles containing large deletions in clones A4 and H3 but not in clones A2 and A12 (Figure 5b). This is probably because the deletion size of the undetected by NGS mutated alleles of A2 and A12 is so large that it causes the removal of the binding site of one of the PCR primers we used for the NGS genotyping (Figure 5a and Supplementary Figure S1).

**Table 2.** Genotyping results of single-cell derived clones. For both CRISPR knockout experiments, the clones are sorted according to the percentage of the wild type allele as determined by NGS. For the ICE analysis, the *Abracl* and *Hprt* clones were Sanger sequenced with the F2A and F1H primers, respectively (see Supplementary Figure S1). In addition, clones A2, A4, A12 and A21 were also sequenced with the R2A primer, while clones H3 and H15 were also sequenced with the F2H primer. Following Synthego’s recommendation of using an R^2^>0.8 for a robust ICE analysis (Synthego, 2019), the genotypes of clones A2, A4, A12 and H3 were classified by both primers as non-determined (R^2^ ≤0.6). The clone A21 was classified as non-determined by the F2A primer. For this clone we present the genotyping result obtained by the R2A primer. Using either the F1H or the F2H primer, the clone H15 was genotyped by ICE as fully mutated (in both cases with the same R^2^ value), in agreement with the results obtained by PIM-RFLP. However, using the Ion Torrent platform, in two independent experiments the wild type allele was detected at a frequency of ∼10% in the H15. Sequencing this sample using the Illumina platform, we confirmed the absence of the wild type allele in this clone. For all other clones, the presenting NGS results were obtained using the Ion Torrent platform. WT: Wild Type, ND: Non-Determined

| *Abrac1* |  |  |  |  |  |  | *Hprt* |  |  |  |  |  |  |
| --- | --- | --- | --- | --- | --- | --- | --- | --- | --- | --- | --- | --- | --- |
| Clone | WT%<br>by<br>NGS | Genotype | | | WT%<br>by<br>ICE | ICE<br>$R^2$ | Clone | WT%<br>by<br>NGS | Genotype | | | WT%<br>by<br>ICE | ICE<br>$R^2$ |
|  |  | NGS | PIM-RFLP | ICE |  |  |  |  | NGS | PIM-RFLP | ICE |  |  |
| A1 | 0 | – | – | – | 0 | 0.88 | H3 | 0 | – | – | ND | ND | 0.10 |
| A2 | 0 | – | – | ND | ND | 0.60 | H9 | 0 | – | – | – | 0 | 0.98 |
| A4 | 0 | – | – | ND | ND | 0.36 | H11 | 0 | – | – | – | 0 | 0.96 |
| A8 | 0 | – | – | – | 0 | 0.94 | H15 | 0 | – | – | – | 0 | 0.97 |
| A9 | 0 | – | – | – | 0 | 0.93 | H23 | 6 | +/- | +/? | +/- | 3 | 0.94 |
| A12 | 0 | – | – | ND | ND | 0.43 | H5 | 12 | +/- | +/- | +/- | 7 | 0.87 |
| A16 | 0 | – | – | – | 0 | 0.94 | H10 | 26 | +/- | +/- | +/- | 20 | 0.94 |
| A21 | 0 | – | – | – | 0 | 0.96 | H17 | 28 | +/- | +/- | +/- | 31 | 0.96 |
| A5 | 8 | +/- | +/- | +/- | 7 | 0.94 | H14 | 36 | +/- | +/? | +/- | 33 | 0.97 |
| A6 | 15 | +/- | +/? | +/- | 15 | 0.95 | H8 | 39 | +/- | +/- | +/- | 38 | 0.96 |
| A7 | 30 | +/- | +/- | +/- | 48 | 0.95 | H12 | 42 | +/- | +/- | +/- | 50 | 0.95 |
| A15 | 31 | +/- | +/- | +/- | 35 | 0.87 | H2 | 50 | +/- | +/- | +/- | 41 | 0.90 |
| A10 | 37 | +/- | +/? | +/- | 42 | 0.95 | H4 | 52 | +/- | +/? | +/- | 48 | 0.98 |
| A14 | 73 | +/- | +/? | +/- | 72 | 0.97 | H7 | 63 | +/- | +/- | +/- | 68 | 0.98 |
| A20 | 75 | +/- | +/? | +/- | 76 | 0.97 | H6 | 65 | +/- | +/? | +/- | 75 | 0.97 |
| A3 | 84 | +/- | +/? | +/- | 81 | 0.98 | H1 | 68 | +/- | +/? | +/- | 69 | 0.98 |
| A18 | 88 | +/- | +/? | +/- | 87 | 0.99 | H16 | 69 | +/- | +/? | +/- | 73 | 0.98 |
| A17 | 93 | +/- | +/? | +/- | 94 | 0.99 | H13 | 70 | +/- | +/? | +/- | 87 | 0.93 |
| A22 | 95 | +/- | +/? | +/- | 96 | 0.99 | H19 | 89 | +/- | +/? | +/- | 86 | 0.99 |
| A13 | 96 | +/- | +/? | + | 100 | 1.00 | H18 | 100 | + | +/? | + | 100 | 1.00 |
| A11 | 97 | +/- | +/? | +/- | 96 | 0.99 | H20 | 100 | + | +/? | + | 100 | 1.00 |
| A19 | 100 | + | +/? | + | 100 | 1.00 | H21 | 100 | + | +/? | + | 99 | 0.99 |
| A23 | 100 | + | +/? | + | 100 | 1.00 | H22 | 100 | + | +/? | + | 100 | 1.00 |

**Figure 5.**
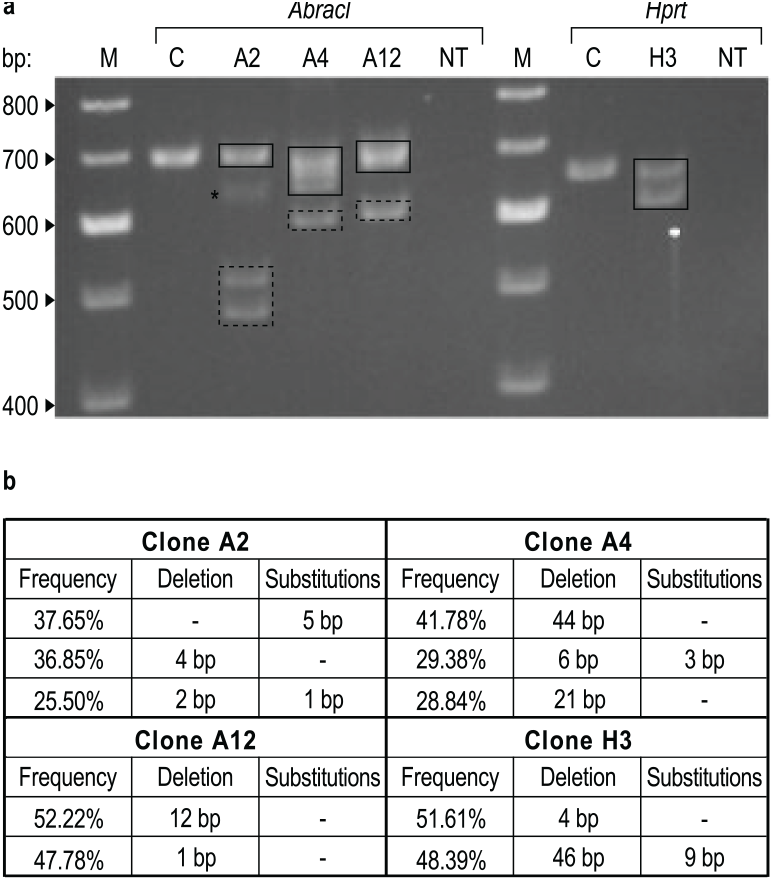
**(a)** Electrophoresis in a 2% (w/v) agarose gel of the PCR products that were Sanger sequenced in an unsuccessful attempt to genotype by ICE the indicated single-cell clones. Unedited Neuro-2a cells were used as control (C). The wild type alleles of the *Abracl* and *Hprt* loci are expected to give a single amplicon of 688 bp and 647 bp, respectively. Gel bands that appears to correspond to alleles identified by NGS are indicated by solid line rectangles. Gel bands within dashed line rectangles correspond to alleles carrying large deletions that makes unlikely or impossible to be PCR amplified with the primers we used for the NGS genotyping, and thus it is not surprising that escaped detection by NGS (see also Supplementary Figure S1). A faint band indicated by a star at the PCR products of clone A2 is about 60 bp smaller than the band produced by the wild type allele. Although the size of this deletion is not large enough to explain why it wasn’t detected by NGS, this allele may also contain a number of substitutions and thus it has lost the binding site of one of the primers, which amplify the 198 bp NGS targeted region. Indeed, by agarose gel electrophoresis of the PCR products produced by clone A2 using the FA/RA pair of primers, we failed to detect any amplicon appreciably smaller than 198 bp (data not shown). **(b)** The allele composition of these clones as determined by NGS. M: DNA marker, NT: Non template control.

## DISCUSSION

As CRISPR-Cas9 becomes a widely embraced system for genome editing, the development of highly accessible (in terms of both speed and cost) and reliable indel detection techniques can help to simplify experimental routines that are currently executed by a vast number of research laboratories. Here we present PIM-RFLP, a new indel detection method, which has the potential to become a valuable tool for the analysis of CRISPR editing results. By comparing PIM-RFLP with NGS we have seen that it is very accurate. Furthermore, it appears to have some advantages (in terms of both sensitivity and speed) over the Sanger sequencing deconvolution methods, which constantly gain popularity because they are cost-effective (like T7E1 assay) and provide information at the allele sequence level (like NGS).

Although we chose to focus on ICE, we also analysed our Sanger sequencing results with TIDE (Brinkman *et al*, 2014) and DECODR (Bloh *et al*, 2021) (data not shown). However, a disadvantage of TIDE over ICE and DECODR is that it is not fully automated and often needs the parameters to be adjusted by the user. Using TIDE, we obtained results very similar to ICE for the *Abracl* data set. TIDE, like ICE, also failed to genotype clones A2, A4 and A12. Analysing the Sanger traces from the *Hprt* locus with TIDE, we also failed to genotype the clone H3 and we consistently obtained lower R^2^ values than with ICE using the F1H primer.

DECODR, in most cases, gave similar R^2^ values as ICE for both loci. Of special interest are the results we obtained with DECODR for the clones A2, A4, A12 and H3 since this algorithm has been designed to allow the genotyping of larger deletions (Bloh *et al*, 2021). Analysing these clones with DECODR we obtained mixed results. DECODR also failed to genotype the clone A12 (R^2^ = 0.72 with the primer F2A, while the analysis with the primer R2A failed), but it gave acceptable R^2^ values (0.86-0.95) for the other three clones. The clones A4 and H3 were correctly genotyped as fully mutated and the sizes of the deletions detected were in line with those observed by agarose gel electrophoresis and NGS (see Figure 5). However, using the F2A primer the clone A2 was genotyped (with R^2^ = 0.89) as fully mutated (in agreement with NGS and PIM-RFLP), but using the R2A primer the wild type allele was incorrectly detected at a frequency of 19% (with R^2^ = 0.86). Furthermore, in clone H7 DECODR failed to detect the presence of the wild type allele (with R^2^ = 0.91).

Concerning the accuracy of PIM-RFLP in the evaluation of the editing efficiency in pools of edited cells, it is not possible to extract safe conclusions due to the limiting size of our data set. However, PIM-RFLP performed better than any decomposition method for both loci we tested. For the analysis of pools by PIM-RFLP, a normaliser gene can be used in order to compensate for technical differences between the edited and the unedited (control) samples. When we analysed the pools of CRISPR-edited cells at the *Abracl* and *Hprt* loci, we used as a normaliser the *Gusb* gene (data not shown). However, we didn’t include the results in the calculation of the editing efficiencies, as both the edited and the control samples gave practically identical Ct values for *Gusb*.

PIM-RFLP is a very flexible method and thus, other variations than the one presented here can be envisioned. Undoubtedly, for performing PIM-RFLP, XcmI is a particularly suitable choice among the rare-cutter restriction enzymes with a degenerate recognition site. The nine unspecific nucleotides of its cleavage site give great flexibility in designing the mutagenic primers. Furthermore, the presence of the dinucleotide GG in its recognition sequence increases the possibility of designing efficient mutagenic primers when working with SpCas9, the most commonly used programmable endonuclease, which recognises the PAM sequence 5′-NGG-3′. However, when designing a PIM-RFLP assay, other enzymes that have a degenerate recognition site should also be considered. Although the mutagenic primers generating the XcmI site worked very efficiently in all the 12 cases we tested, for three genes, choosing a different enzyme we could have designed primers with fewer mismatches and/or with mismatches further away from the 3′ end. In *Gusb*, for example, our forward mutagenic primer contains a mismatch at position −3, whereas the reverse contains three mismatches at positions −6 to −8. Using BglI, which recognises the sequence 5′-GCCN_5_GGC-3′, it is possible to design mutagenic primers with only two mismatches, one at position −6 of the forward primer and another at position −3 of the reverse.

Another possible modification could be to change the location of the mutagenic primers relative to the DSB. For example, the 3′ end of the primer can be moved one nucleotide closer to the DSB. However, the most common mutated alleles following CRISPR-Cas9 targeting are those with 1 bp insertion or deletion (Chakrabarti *et al*, 2019; Chen *et al*, 2019; Allen *et al*, 2019). Our design allows the amplification of alleles carrying 1 bp deletions (or insertions) while it makes improbable the amplification of alleles carrying 3 bp (in frame) deletions. Therefore, when single cell clones are genotyped, a PCR product that is completely indigestible indicates the presence of a fully mutated clone, carrying at least one targeted allele that is highly probable to contain a frameshift, loss of function, mutation. Such a clone is more likely to be fully knocked out for the target gene than a clone for which the only available information is that it carries exclusively non-amplifiable mutated alleles.

Apparently, PIM-RFLP is not free of shortcomings. An unusual prerequisite of our method is that the mutagenic primers must be PAGE purified. In preliminary experiments using standard High-Performance Liquid Chromatography (HPLC) purified primers, about 20% of the PCR product produced by the wild type allele remained uncut after the XcmI digest. Apparently, the presence of shorter versions of the mutagenic oligonucleotide primers resulted in amplicons carrying the indigestible sequence 5′-CCAN_<9_TGG-3′. As it can be seen in Figure 2, this problem was not completely solved by using PAGE purified primers. In few cases, a faint ghost indigestible band remains visible after the XcmI treatment. Probably this is because of suboptimal purification of at least one of the two mutagenic primers that accomplish the amplification. A faint band also appeared in some of the *Abracl* and *Hprt* clones we examined (see Figure 4). Fortunately, such ghost bands do not compromise the ability of PIM-RFLP to reliably identify fully mutated clones. Furthermore, when it comes to the evaluation of the editing efficiency in pools of cells, using as control the digest of the PCR product from the corresponding unedited cells, necessary adjustments can be done to obtain reliable results.

When compared with the Sanger sequencing deconvolution methods, PIM-RFLP has the main disadvantage of not providing information at the sequence level for the mutated alleles. However, it also has some important advantages. Firstly, it performs better in genotyping fully mutated clones that carry large deletions. ICE failed to genotype 3 out of 8 fully mutated clones for the *Abracl* and 1 out of 4 fully mutated clones for the *Hprt*. We have seen that the presence of even a single allele that carries a large deletion can highly compromise the ability of the Sanger sequencing deconvolution methods to provide reliable genotyping results. This is a potentially serious shortcoming, especially when working with polyploid cancer cell lines or polyploid plant cells, since the possibility of having at least one allele carrying a large deletion increases. Secondly, PIM-RFLP can be performed in a single day using standard molecular biology laboratory equipment. This is important for laboratories that do not have in-house Sanger sequencing services. Thirdly, like any RFLP assay, it is a very simple and intuitive method. This makes troubleshooting easier and diminishes the possibility of accepting erroneous results.

Although NGS remains the most reliable allele profiling method, its high cost and extended turnaround time from samples to results limits its use for assessing the outcome of CRISPR gene knockout experiments. As a much more accessible alternative, T7E1 assay has been widely used because of its low instrumentation requirements, cost-effectiveness and fast turnaround time. The T7E1 assay has been historically proven to be a valuable technique and it is still used as a fast and economical first test for analysing the results of CRISPR knockout experiments. However, it can produce erroneous results when it is used either to assess the editing efficiency in pools of edited cells (Sentmanat *et al*, 2018) or to identify cell clones that carry exclusively mutated alleles (Yang *et al*, 2015). ICE and other sequencing deconvolution algorithms are currently the most popular choices for the detection of CRISPR-induced indels. However, their ability to genotype clones carrying large deletion alleles is limited, their turnaround time can be relatively long, and although they are reliable, the potentially fully mutated clones must be further validated (Synthego, 2022) using NGS, digital PCR (Mock *et al*, 2016) or a method that confirms the absence of the target gene’s product. In this landscape, we believe that PIM-RFLP can be a very useful addition to the toolbox of CRISPR genotyping methods. It compares favourably with T7E1 assay since it is equally accessible, cost-effective and fast but much more reliable. Thus, it can replace the T7E1 assay as a first test in the analysis of CRISPR experimental results, allowing the shortlisting of fully mutated, and potentially fully knockout, single-cell clones (including clones with large deletion alleles that tend to compromise the analysis by Sanger sequencing deconvolution algorithms). Promising clones can be then independently validated by other methods, including Western blot or any assay that assesses the function of the target gene’s product. For laboratories that do not have direct access to Sanger sequencing facilities, this approach enables the identification of fully knocked out clones using exclusively in-house techniques, facilitating further the analysis of CRISPR editing results. It has been noticed that one of the revolutionary properties of CRISPR methodology is that it made gene editing highly accessible, effectively “democratising” the technology (Ledford, 2015; Jasin & Haber, 2016). We believe that PIM-RFLP will further facilitate CRISPR’s accessibility by providing a simple, fast and economical indel detection option.

## ACKNOWLEDGEMENTS

We would like to thank Anagnostis Argiriou and George Tsiolas from the Institute of Applied Biosciences / CERTH (Thessaloniki, Greece), who conducted the Illumina NGS of clone H15.

