## Supplementary Data for "A PCR-RFLP method for the detection of CRISPR-induced indels"

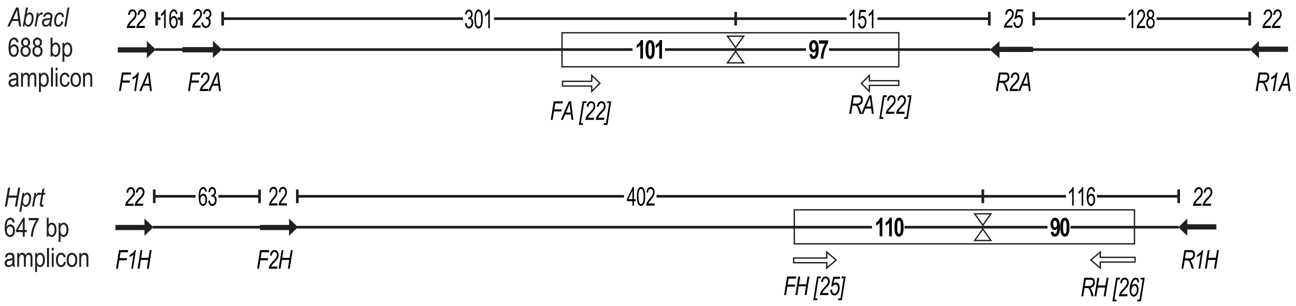


**Supplementary Figure S1.** Schematic representation of the PCR amplicons that were Sanger sequenced for ICE genotyping. The primers used for the PCR amplification and/or the subsequent Sanger sequencing are indicated by black arrows. The length and the name of them are given in italics above and below the arrows, respectively. A pair of open arrowheads facing each other marks the position of the DSB. The plain numbers denote distances between primers or between a primer and the DSB. All the 688 bp *Abracl* amplicons were sequenced with the F2A primer. The amplicons obtained by clones A2, A4, A12, and A21 were also sequenced with the R2A primer. All the 647 bp *Hprt* amplicons were sequenced with the F1H primer. The amplicons obtained by clones H3 and H15 were also sequenced with the F2H primer. Rectangles outline the regions targeted by NGS after PCR amplification using the pairs of primers FA/RA and FH/RH (indicated by white arrows with their length given in brackets) for the *Abracl* and the *Hprt* loci, respectively. By bold numbers inside the rectangles is given the length of the NGS-determined sequences flanking the DSB. Mutated alleles carrying large deletion extending more than about 65-85 bp to the same direction of the DSB escape detection by NGS since they fail to PCR amplify with the primers we used for the NGS genotyping.

| Gene | gRNA target (5′ to 3′) | NCBI Accession No |
|---|---|---|
| *Abracl* | GTGGAGGAAATTCATCGCCT | NM_028440 |
| *Hprt* | ACCTCTTAGGAGTCTAAAGT | NM_013556 |

**Supplementary Table S1.** The gRNA targets of the *Abracl* and *Hprt* genes. The *Abracl* gRNA was designed using the Custom Alt-R CRISPR-Cas9 guide RNA online tool provided by IDT (https://www.idtdna.com/site/order/designtool/index/CRISPR_CUSTOM). The *Hprt* gRNA was included in the mouse Alt-R CRISPR-Cas9 Control Kit (IDT).

| Mutagenic primers (supplied by IDT) | | | |
|---|---|---|---|
| Name | Sequence 5´ to 3´ | Length | Tm |
| F_Abracl | CATGAAGTTAACCTCCTGGTGGAGGccATTCATC | 34 | 65 |
| R_Abracl | ATTCGGCCTTAGAGTACTTACTTCTGGAACCCAG | 34 | 64 |
| F_Agpat1 | AGGTGCGCGGGGCTCACCACTTCCCTCCCAC | 31 | 74 |
| R_Agpat1 | AGGGAACTCTGGTGGTTGGACACAACCACcaAGGGCT | 37 | 71 |
| F_Canx | CACCTTTGTCTCTGAAGGATCAATTCCACG | 30 | 61 |
| R_Canx | CACTTATCTGGACCAAACATAATAGTATAGccAGTCTT | 38 | 61 |
| F_Capn2 | CCCGAGCCAGGGAGCGGTCAGATACCTTCAccAACCTC | 38 | 72 |
| R_Capn2 | GGGGGCAGCTTGAAGCGGTTGAGGACCaCG | 30 | 71 |
| F_Ctbp1 | CACACGTCTGCCCTCCTCTCTCAGCAGGTACCaTCC | 36 | 70 |
| R_Ctbp1 | TGGGATGCCAGTGGGGGCCACACTCcCaACGC | 32 | 74 |
| F_Eif2b2 | GATGACGGCTGCGCAGCCCTCCaAGACCA | 29 | 71 |
| R_Eif2b2 | CCGGATGATCTTGAGGACTCTCCGCACCATATTTCCCAC | 39 | 69 |
| F_Gusb | AACAACACACTGACCCCTCATACCCTTCCaCC | 32 | 66 |
| R_Gusb | CTCACATGGAGGTGTCAGTCTTGTAGACccaGGTCC | 36 | 67 |
| F_Hmbs | GACCGCAGCGGCACTGCCGCGCCTGCGCCaTG | 32 | 79 |
| R_Hmbs | GTTGTGGCCGCGCCGCCGTTACCGGACAccACTCCG | 36 | 77 |
| F_Hprt | TAATTTATGCCTGCAACCCTACCTCTTAccAGTCTA | 36 | 63 |
| R_Hprt | GGTTGGTGGCCTTAAACTCAAGATCCCCCaAC | 32 | 66 |
| F_Tbp | GCTGCAGTCATCATGAGAATAAGAGAGCCACG | 32 | 64 |
| R_Tbp | ACACCATTTTTCCAGAACTGAAAATCAACcaAGTTG | 36 | 62 |
| F_Tfrc | GCTTTGGGTGCTGGTGTTGCGGCcAAGTCC | 30 | 70 |
| R_Tfrc | ATACTTGGGCAAGTTTCAACAGAAGACCTGTTCCCACA | 38 | 66 |
| F_Trim27 | GTGCCTGGCCCGCTGCTGGGGTGCCaCG | 28 | 76 |
| R_Trim27 | CGGGAAGGTCTCCCGGCACTGCGGACACGACACcaTAGTC | 40 | 74 |
| Primers used for the NGS analysis (supplied by Eurofins Genomics) | | | |
| Name | Sequence 5´ to 3´ | Length | Tm |
| FA | TGCTGGGAGTACAAAATGGCTG | 22 | 57 |
| RA | ACTGTCCTCAGTCCTAAGTGGC | 22 | 58 |
| FH | GTTTGGGATGTTAAGAGTCCCTATC | 25 | 55 |
| RH | GCACACATACACAAATCTTTCTGTTG | 26 | 56 |
| Primers used for the ICE analysis (supplied by Eurofins Genomics or IDT) | | | |
| Name | Sequence 5´ to 3´ | Length | Tm |
| F1A | TCCTGTTACCTTTTGCAGTTTT | 22 | 53 |
| F2A | CTGACACAAGCTGTGCATGGAGT | 23 | 60 |
| R1A | CCATGATTTTAACAACAAACCA | 22 | 50 |
| R2A | ACTCCCCAATAGCCCTATGTCTCAG | 25 | 59 |
| F1H | AGGTTTCGAGCCCTGATATTCG | 22 | 57 |
| F2H | TCTAGGGCATAATGGGTGCCAT | 22 | 58 |
| R1H | ATGTGGCAAGGTCAAAAACAGT | 22 | 56 |

**Supplementary Table S2.** The primers used in this study. Forward primers start with F, while reverse primers start with R. The melting temperatures (Tm) were calculated using AmplifX version 2 (by Nicolas Jullien; Aix-Marseille University, CNRS, INP, Institute of NeuroPhysiopathology, Marseille, France - https://inp.univ-amu.fr/en/amplifx-manage-test-and-design-your-primers-for-pcr). The mismatches between the mutagenic primers and their templates are shown in lowercase.

| Gene | gRNA target (5´ to 3´) | NCBI Accession No |
|---|---|---|
| *Agpat1* | ACAACCACGTAGGGCTGCGT | NM_018862 |
| *Canx* | TAGTATAGGGAGTCTTGTCG | NM_001110500 |
| *Capn2* | TACCTTCATTAACCTCCGCG | NM_009794 |
| *Ctbp1* | GCCACACTCACGACGCCTGG | NM_001198860 |
| *Eif2b2* | CAGCCCTCCGAGACCACCGT | NM_145445 |
| *Gusb* | TTGTAGACGATGGTCCCCGG | NM_010368 |
| *Hmbs* | ACCGGACATGACTCCGCCCA | NM_013551 |
| *Tbp* | AAAATCAACGCAGTTGTCCG | NM_013684 |
| *Tfrc* | TGTTGCGGCGAAGTCCAGTG | NM_011638 |
| *Trim27* | ACACGACACGTTAGTCTCCG | NM_009054 |

**Supplementary Table S3.** The gRNA targets suggested from the CHOPCHOP web tool for the ten randomly selected housekeeping genes.
